# Motor skill learning decreases motor variability and increases planning horizon

**DOI:** 10.1101/505198

**Authors:** Luke Bashford, Dmitry Kobak, Jörn Diedrichsen, Carsten Mehring

## Abstract

We investigated motor skill learning using a path tracking task, where human subjects had to track various curved paths at a constant speed while maintaining the cursor within the path width. Subjects’ accuracy increased with practice, even when tracking novel untrained paths. Using a “searchlight” paradigm, where only a short segment of the path ahead of the cursor was shown, we found that subjects with a higher tracking skill differed from the novice subjects in two respects. First, they had lower motor variability, in agreement with previous findings. Second, they took a longer section of the future path into account when performing the task, i.e. had a longer planning horizon. We estimate that between one third and one half of the performance increase was due to the increase in planning horizon. An optimal control model with a fixed horizon (receding horizon control) that increases with tracking skill quantitatively captured the subjects’ movement behaviour. These findings demonstrate that human subjects not only increase their motor acuity but also their planning horizon when acquiring a motor skill.

**New and Noteworthy:** We show that when learning a motor skill humans are using information about the environment from an increasingly longer amount of the movement path ahead to improve performance. Crucial features of the behavioural performance can be captured by modelling the behavioural data with a receding horizon optimal control model.

## Introduction

The human motor system can acquire a remarkable array of motor skills. Informally, a person is said to be “skilled” if he or she can perform faster and at the same time more accurate movements than other, unskilled, individuals. What we don’t know, however, is what learning processes and components underlie our ability to move better and faster. One component may be relatively “cognitive”, involving the faster and more appropriate selection and planning of upcoming actions (Diedrichsen and Kornysheva 2015; Wong et al. 2015). Another component may be related to motor execution – the ability to produce and fine control difficult combinations of muscle activations (Shmuelof et al. 2012; Waters-Metenier et al. 2014). Depending on the structure of the task, changes in visuo-motor processing or feedback control may also contribute to skill development. Motor adaptation extensively studied using visuomotor and force perturbations (Shadmehr et al. 2010), may play a certain role in stabilizing performance, but it cannot by itself lead to improvements in the speed-accuracy trade-off (Wolpert et al. 2011).

A task commonly used in the experiments on motor skill learning is sequential finger tapping, where subjects are asked to repeat a certain tapping sequence as fast and as accurately as possible (Karni et al. 1995, 1998; Petersen et al. 1998; Walker et al. 2002). Improvement in such a task can continue over days, but previous papers have focussed mostly on the learning that is specific to the trained sequence(s) (Karni et al. 1995).

Many real-world tasks, however, do not involve the production of a fixed sequence of motor commands, but the flexible planning and execution of movements. Such flexibility is often well described by optimal feedback control models (Braun et al. 2009; Diedrichsen et al. 2010; Todorov and Jordan 2002) where the skilled actor appears to compute “on the fly” the most appropriate motor command for the task at hand. This requires demanding computations (Todorov and Jordan 2002), and the human motor system likely has found heuristics to deal with this complexity. One way to reduce complexity of the control problem is to not optimize the whole sequence of motor commands that will achieve the ultimate goal, but to only optimise the current motor command for a short distance into the future. This idea is called receding horizon control, also known as model predictive control (Kwon and Han 2005). Under this control regime, the system computes a feedback control policy that is optimal for a finite planning horizon. The control policy is then continuously updated as the movement goes on and the planning horizon is being shifted forward. This allows for adaptability, e.g. it can flexibly react to perturbations or unexpected challenges, as sensory information becomes available. Recent studies provided indirect evidence that favour the optimisation of short time-periods of a motor command (Dimitriou et al. 2013). The notion of planning horizon also arises in reinforcement learning, e.g. in the context of the so-called successor representation (Momennejad et al. 2017).

Motivated by these ideas, we propose that some of the skill of a down-hill skier or a race-car driver may lie not only in the increased ability to execute difficult motor commands (e.g. due to decreased motor variability), but also in the ability to plan further ahead and to optimize the movements for a longer time period into the future. In addition, we propose that the time span that subjects plan ahead increases with experience, leading to an increasing performance with training.

To test this idea, we designed an experimental condition which would allow us to measure the planning horizon that skilled actors are using when executing long sequence of movements that need to be planned “on the fly” – i.e. where the actual sequence of movements cannot be memorized. For this, we developed a path tracking task, where subjects had to maintain their cursor within a path that was moving towards them at a fixed speed. A similar task has been previously used in motor control research (Poulton 1974), using a mechanical apparatus with paths drawn on a paper roll that was moving at a fixed speed. It has been shown that subjects are able to increase their accuracy with training, but the different computational strategies between expert subjects and naïve performers remain unclear. In our study we use ‘searchlight’ trials in which subjects see various lengths of the approaching path ahead of their cursor to probe subjects forward planning and compare experts and novices in this respect.

## Materials and Methods

### Subjects

62 experimentally naïve subjects took part in this experiment (33 males and 29 females, age range 20-52 years old). Subjects gave written informed consent and were paid 10 €/h. The experimental procedures received ethics approval from the University of Freiburg.

### Setup

Subjects sat at a desk looking at a computer monitor (Samsung Syncmaster 226BW) located ~80cm away. A cursor displayed on the screen (Matlab and Psychophysics Toolbox Version 3 (Brainard 1997)) was under position control by movements of a computer mouse. The mouse could be moved on the desk in all directions but only the horizontal (left and right) component contributed to the cursor movement: the vertical position of the cursor was fixed at 5.7mm above the base of the screen.

### Task

To begin each trial subjects had to press the space bar. This displayed the cursor (R=2.9mm, 1.1cm from the bottom of the screen) and the path (width = 2.83cm) that extended from the top to bottom of the screen (30cm). The path continuously moved downward on the screen at a vertical speed of 34.1cm/s. The initially visible path was a straight line centered in the middle of the screen with the cursor positioned in the middle of the path. Once this initial section moved through the screen, the path then followed a random curvature (Fig. 1A). Subjects were instructed to keep the cursor between the path borders at all times moving only in the horizontal plane and were told to be as accurate as possible. The cursor and path were displayed in white if the cursor was within the path and both turned red when it was outside the path, always on a black background.

**Figure 1.**
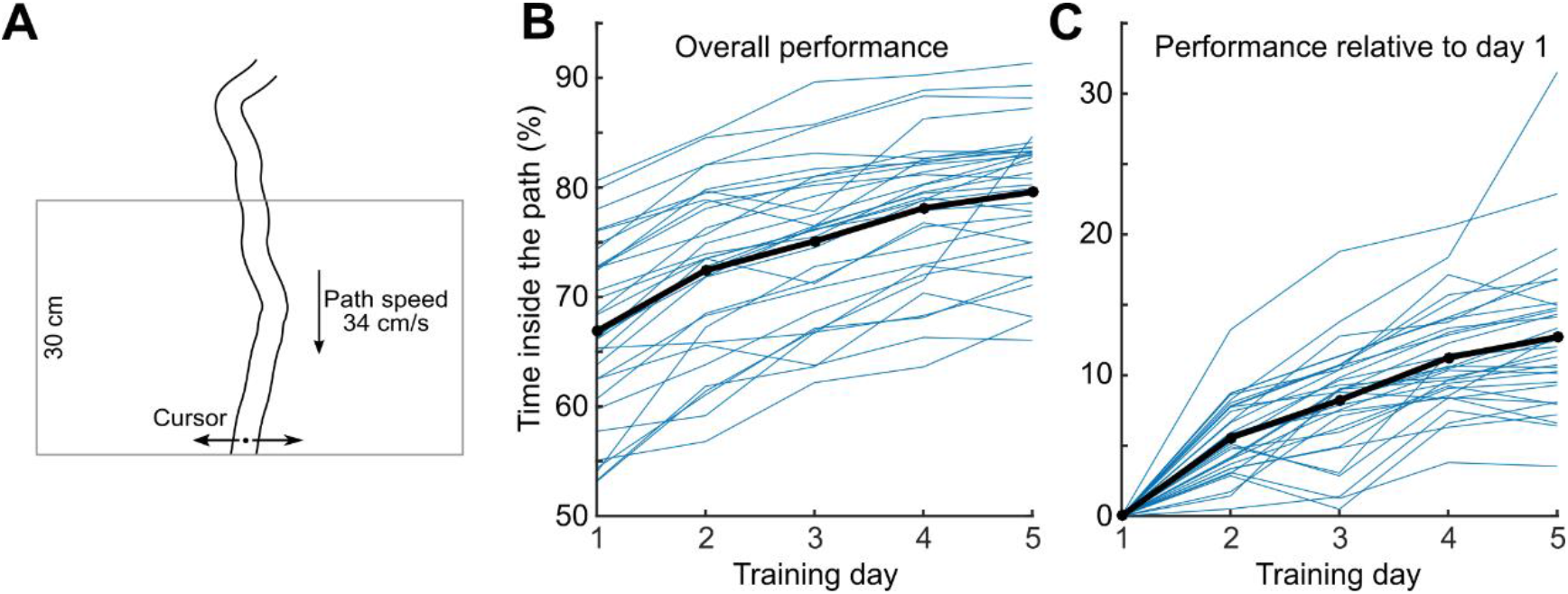
Experimental Paradigm. (A) Subjects had to track a curved path that was dropping down from top to bottom of the screen with a fixed speed of 34 cm/sec by moving the cursor horizontally. (B) Expert subjects’ performance over the 5 days of training. Bold line shows the group average, thin lines show individual subjects (each point is a mean over 3 trials with the same searchlight length, 100%). (C) Expert subjects’ performance over the 5 days of training with the performance on the first day subtracted.

The cursor position was sampled at 60 Hz and the tracking accuracy was defined for each trial as the percentage of time steps when the cursor was inside the path. Running accuracy values were continuously displayed in the top left corner of the screen and final accuracies were displayed between the trials.

This experiment is based on a previous version where subjects were asked to track static randomly curved paths in 2D as quickly as possible without touching the sides [unpublished data, (Bashford et al. 2014)]. We later found that the 1D paradigm presented here was better suited to study the planning horizon as the speed was fixed.

### Paradigm

Subjects were randomly assigned into two groups: expert (N=32) and naive (N=30). The paradigm included a training (expert group only) and a testing (all subjects) phase. Subjects in the expert group trained over 5 consecutive days, each day completing 30 minutes of path tracking (10 of 3-minute trials with short breaks in-between, searchlight length (s) 100%). If the performance improved from one trial to the next subjects saw a message saying “Congratulations! You got better! Keep it up!”, otherwise the message “You were worse this time! Try to beat your score!” was shown. The training paths were randomly generated on the fly. Experts performed the testing set of trials after a short break following training on the final (5th) day. Naïve subjects performed only the testing set of trials.

The testing phase lasted 30 min (30 of 1-minute trials with breaks in-between) using 30 different pre-generated paths that were the same for all subjects. The testing phase in this experiment contained 3 normal trials (s=100%) and 27 searchlight trials (s=10-90%) where some upper part of the path was not visible. Three blocks of 10 trials with the searchlight length ranging from s=10% to s=100% (in steps of 10%) were presented, with the order shuffled in each block; the same fixed pseudorandom sequence was used for all subjects.

### Path generation

Paths were generated before each trial start during training and a pre-generated fixed set was produced in the same way for testing. Each path was initialized to start at the bottom middle of the screen and the initial 30 cm of each path were following a straight vertical line. Subsequent points of the path midline had a fixed Y step of 40 pixels (1.1 cm) and random independent and identically distributed (iid) X steps drawn from a uniform distribution from 1 to 80 pixels (2.7mm – 2.2cm). Any step that would cause the path to go beyond the right or left screen edges was recalculated. The midline was then smoothed with a Savitzky-Golay filter (12th order, window size 41) and used to display path boundaries throughout the trial. All of the above parameters were determined in pilot experiments to create paths which were very hard but not impossible to complete after training.

### Statistical analysis

In all cases, we used nonparametric rank-based statistical tests to avoid relying on the normality assumption. In particular, we used Spearman’s correlation coefficient instead of the Pearson’s coefficient, Wilcoxon signed-rank test instead of paired two-sample t-test, and Wilcoxon-Mann-Whitney ranksum test instead of unpaired two-sample t-test.

We initially recorded N=10 subjects in each group and observed statistically significant (p<0.05) effect that we are reporting here: positive correlation between the asymptote performance and the horizon length, as estimated via the changepoint and exponential models. We then recorded another N=20/22 (naïve/expert) subjects per group to confirm this finding. This internal replication confirmed the effect (p<0.05). The final analysis reported in this study was based on all N=62 subjects together. A preliminary version of the analysis for the initial N=10/10 subjects can be found in our preprint (Bashford et al. 2014), but note that it used a different way to estimate planning horizon compared to the procedure presented here, and so the values are not directly comparable.

### Changepoint and Exponential model

We used two alternative models to describe the relationship between the searchlight length and the accuracy: a linear changepoint model and an exponential model. We used two different models to increase the robustness of our analysis and both models support our conclusions.

The changepoint model is defined by

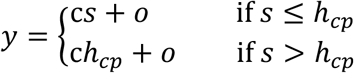

where *y* is the subject’s performance, *s* the searchlight length and (*c*, *o*, *h*_cp_) are the subject-specific parameters of the model which define the baseline performance at searchlight 0% (*o*), the amount of increase of performance with increasing searchlight (*c*) and the planning horizon (*h*_cp_) after which the performance does not increase any further.

The exponential model is defined by

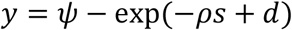

where the subject-specific parameters (*ψ*, *d*, *ρ*) specify the performance at searchlight 0% (*φ* − exp[*d*]), the asymptote for large searchlights (*ψ*) and the speed of performance increase (*ρ*). This function monotonically increases but it never plateaus. The speed of the increase depends on the parameter *ρ* with larger values meaning faster approaching the asymptote. We used the following quantity as a proxy for the “effective” planning horizon: 10+log(5)/*ρ*. It can be understood as the searchlight length that leads to performance being five times closer to the asymptote than at s=10%. The log(5) factor was chosen to yield horizon values of roughly the same scale as with the changepoint model above.

Both models (changepoint and exponential) were fit to the raw performance data of each subject, i.e. to the 30 data points, 3 for each of the 10 searchlight length values. The exponential fit (see Equation 2 in the Results) was done with the Matlab’s nlinfit() function, implementing Levenberg-Marquardt nonlinear least squares algorithm. The changepoint fit (see Equation 1 in the Results) was done with a custom script that worked as follows. It tried all values of *h*_cp_ on a grid that included s=10% and then went from s=20% to s=100% in 100 regular steps. For each value of *h*_cp_ the other two parameters can be found via linear regression after replacing all *s*>*h*_cp_ values with *h*_cp_. We then chose *h*_cp_ that led to the smallest squared error.

### Trajectory analysis

To shed light on the learning process we analysed additional parameters of the subjects’ movement trajectories.

First, we computed the time lag between the subjects’ movement trajectories and the midline of the paths (Figure 4A-B). To compute the lags, we interpolated both cursor trajectories and path midlines 10-fold (to increase the resolution of our lag estimates). We computed the Pearson correlation coefficient between cursor trajectory and path midline for time shifts from of −300 to 300 ms, and defined the time lag as the time shift maximizing the correlation. Second, we extracted the cursor trajectories in all sections across all paths that shared a similar curved shape to explore the differences in cursor position at the apex of the curve (Figure 4C). The segments were selected automatically by sliding a window of length 18 cm across the path. We included all segments that were lying entirely to one side (left or right) of the point in the middle of the sliding window (“C-shaped” segments), with the upper part and the lower part both going at least 4.5 cm away in the lateral direction (see Figure 3). Our results were not sensitive to modifying the exact inclusion criteria.

To draw the 75% coverage areas of the path inflection points in each group (Figure 4C), we first performed a kernel density estimate of these points using the Matlab function kde2d(), which implements an adaptive algorithm suggested in (Botev et al. 2010). After obtaining the 2d probability density function p(x), we found the largest h such that ∫p(x)dx>0.75 over the area where p(x)>h. We then used Matlab’s contour() function to draw contour lines of height h in the p(x) function.

### Receding horizon model

We modelled subjects’ behaviour by a stochastic receding horizon model in discrete time *t*. In receding horizon control (RHC,(Kwon and Han 2005)) motor commands *u*_*t*_ are computed to minimize a cost function *L*_*t*_ over a finite time horizon of length *h*:

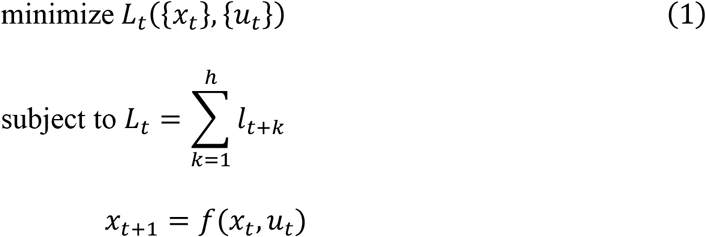

where *f* defines the dynamics of the controlled system. Equation (1) is equivalent to an optimal control problem over the fixed future interval [*t* + 1, *t* + ℎ]. Solving (1) yields a sequence of optimal motor commands 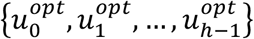. The control applied at time *t* is the first element of this sequence, i.e. 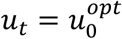. Then, the new state of the system *x*_*t*+1_ is measured (or estimated) and the above optimization procedure is repeated, this time over the future interval [*t* + 2, *t* + 1 + ℎ], starting from the state *x*_*t*+1_.

Applying RHC to our experimental task, the dynamics of the cursor movement was modelled by a linear first-order difference equation:

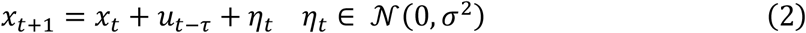

where *t* is the time step, *x*_*t*_ the cursor position at time *t*, *u*_*t*_ is the motor command applied at time *t* and *τ* the motor delay. *η*_*t*_ is the motor noise which was modelled as additive Gaussian white noise with zero mean and variance *σ*^2^. We used the following cost function

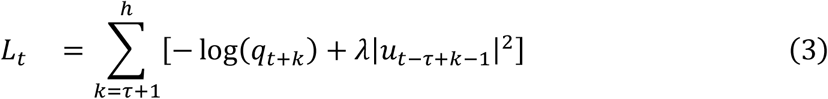

where *L*_*t*_ is the expected cost at time *t*, *q*_*t*+*k*_ is the probability of the cursor being inside the path at time *t+k*, *h* is the length of the horizon in time and *λ* is the weight of the motor command penalty. At every time step *t*, *L*_*t*_ is minimized to compute *u*_*t*_ while {*u*_0_, …, *u*_*t*−1_} are known. Consequently, the lower bound of the sum in (3) is *τ* + 1. The cost function in (3) reflects a trade-off between accuracy (first term, i.e. log[*qt*+*k*]) and effort (second term) whereas their relative importance is controlled by *λ*. Cost functions with a similar accuracy-effort trade-off have been used previously to successfully model human motor behaviour (Braun et al. 2009; Diedrichsen 2007; Todorov and Jordan 2002).

We assume that subjects have acquired a forward model of the control problem and they can, therefore, predict the cursor position at time *t*+1 from the cursor position at time *t* and the motor command in accordance with equation (2). We also assume that subjects have an accurate estimate of the position of the cursor at time *t*, i.e. *x*_*t*_ is known. Subjects can then compute the probability distribution of the cursor position at future times *t*+*k,* given by:

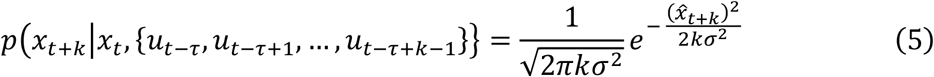

with

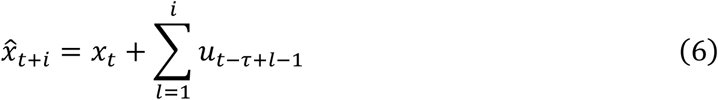

The probability of the cursor being inside the path is then given by

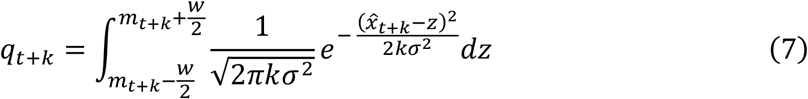

where *m*_*t*_ is the position of the midline of the path at time *t* and *w* the width of the path. The receding horizon model assumes that motor commands *u*_*t*_ are computed by minimizing the cost *L*_*t*_ in each time step *t* for a fixed and known set of model parameters (ℎ, *λ*, *τ*, *σ*^2^). We simplify the optimisation problem by approximating *q*_*t*+*k*_ by

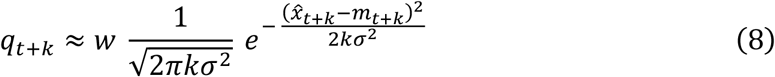

The higher 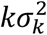 is relative to the path width *w*, the higher the accuracy of this approximation. Note that the squared error is scaled by *kσ*^2^ and hence, errors in the future are discounted. This is a consequence of the used model of the cursor dynamics in (equation 2).

Using equation (8) and removing all terms which do not depend on *u*_*t*_, we can derive a simplified cost function

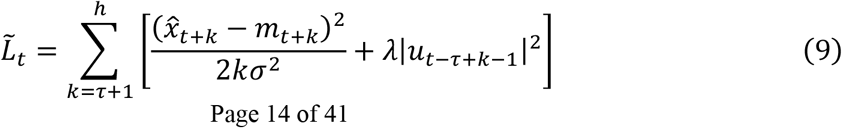

Equation (9) shows that the trade-off between accuracy and the magnitude of the motor commands is controlled by *σ*^2^*λ*. We therefore can eliminate one parameter and use the equivalent cost function

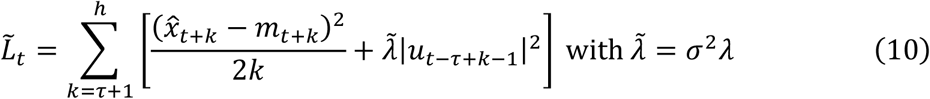

The gradient of the cost function 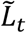 is given by

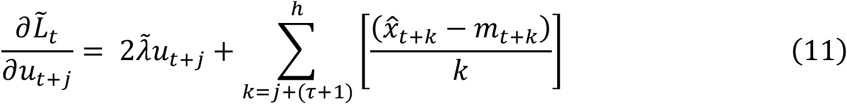

with *j* = 0, …, ℎ − (*τ* + 1). The Hessian of the cost function is given by

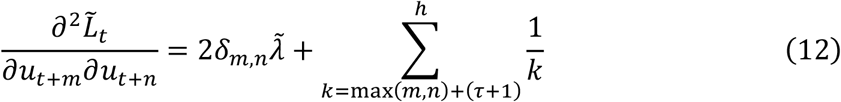

with m, n = 0, …, ℎ − (*τ* + 1). For 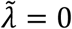 all pivots of the Hessian matrix are positive and therefore the Hessian is positive definite for 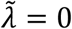. For the general case 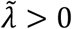 the Hessian remains positive definite as *H*_2_ = *H*_1_ + *D* is positive definite if *H*_1_ is positive definite and *D* is a diagonal matrix with only positive diagonal entries. Given the positive definiteness of the Hessian we can conclude that the cost function 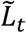 is strictly convex with a unique global minimum. Setting the gradient (12) to **0** defines a system of *h*−*τ* linear equations with *h*−*τ* unknowns (*u*_*t*_, …, *u*_*t*+ℎ−(*τ*+1)_) which solution minimizes 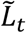. The solution can be computed efficiently using standard numerical techniques. We used the ‘linsolve’ function of MATLAB (R2016b) which uses LU factorization.

As a measure of task performance, we computed the expected time inside the path from the model trajectory *z*_*t*_ as follows

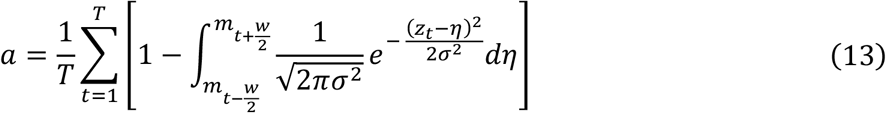

with *T* depicting the number of time steps per path. The lag was computed by maximizing the correlation coefficient between the model trajectories and the path midline identical to how the lag was computed for the subjects’ trajectories.

When applying the model to the searchlight path we made the additional assumption that the model horizon increases with searchlight length *s* up to a maximal value ℎ_*max*_ beyond which the model horizon remains constant:

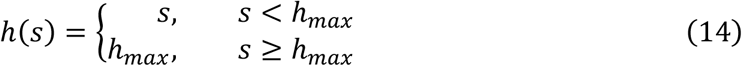

### Fitting the receding horizon model to subjects’ behaviour

We fitted the RHC model to the subjects’ movement trajectories in the searchlight testing paths using Bayesian inference (Gelman et al. 2003). The model parameters were estimated by computing their expected values from the posterior distribution

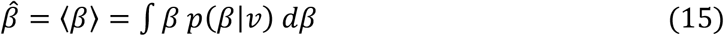

where *β* is the model parameter, *v* the movement trajectory data of a subject and *p*(*β*|*v*) the posterior probability distribution for *β*. We approximated the integral in (15) by sampling from the posterior distribution using the Metropolis algorithm which can sample from a target distribution that can be computed up to a normalizing constant (Gelman et al. 2003). The RHC model has four parameters 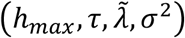 out of which three 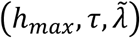 affect the shape of the trajectory (cf. equation (10)). Assuming a flat prior for the model parameters, i.e. . 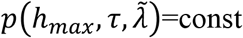., and a non-informative prior for the error-variance *δ*^2^, i.e. *p*(*δ*^2^) = 1⁄*δ*^2^ (Gelman et al. 2003), we obtained the following equation for the posterior

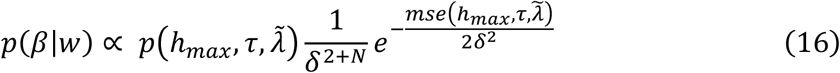

where 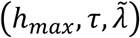 is the mean squared error between the model and the subject movement trajectories and *N* the number of trials. The mean squared error between the movement trajectories of a subject and the model is given by

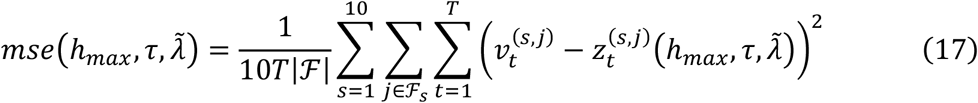

with *T* depicting the number of time steps per path, ℱ_*s*_ the set of paths ids for searchlight *s*, 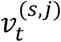 the movement of subject *i* at time *t* in path *j* for searchlight *s* and 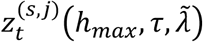 the corresponding movement predicted by the RHC model.

To save computation time, we precomputed the *h*_*max*_ for specific discrete combinations of the model parameters. The model horizon parameter *h*_*max*_ could take any integer value between 1 and 26 given a maximum possible planning horizon of 30cm (vertical screen size) which is equivalent to 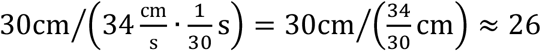 time steps, where 34 cm/s is the path speed and 1/30s the time step. Hence, admissible values for the horizon parameter corresponded to horizons of 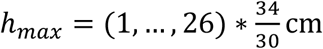. For the delay we allowed the values 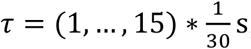, assuming that subjects won’t have larger delays than 500ms. In fact, the maximum delay of a subject we found from fitting was 286 ms which is well below the limit we imposed. The motor penalty parameter 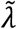 was allowed to take any of 10^3^ logarithmically equally spaced values between 10^−4^ and 10^7^ and 0. In total, we had, therefore, 26×15×1001=390390 admissible parameter combinations for ℎ_*max*_, *τ* and 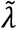. We simulated the model for all of these parameter values and computed the mean squared errors according to equation (17). We then used the Metropolis algorithm to generate 10^6^ samples from the posterior distribution of the parameters. Each sample consisted of a 4-tuple of values for the parameters 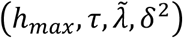. We computed the motor noise parameter of the model *σ*^2^ from the estimated error-variance *δ*^2^ as explained below and then 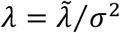 (cf. equation 10). For each parameter sample we also computed the lag, as explained at the end of the previous section, and the task performance using equation (13). As a result, we obtained 10^6^ parameter values, lags and task performances, which reflect samples from the posterior distribution of the model parameters.

To evaluate the quality of the model, we used three-fold cross-validation where in each fold the posterior distributions of the model parameters were estimated using the data from two of the three trials for each searchlight. The posterior distributions were then used to make model predictions of performance and lag in the remaining trial for each searchlight. This was done for each subject separately and the model predictions were compared to the experimentally observed performances and lags (cf. Fig. 5A-D).

Expected values of the model parameters were computed according to equation (13). Expected values were calculated for each cross-validation fold separately and then averaged across the three cross-validation folds. This yielded the model parameters *h*_*max*_, *τ*, *λ*, *σ*^2^ for each subject, shown in Fig. 5E-H.

#### Estimation of the motor noise parameter from the error-variance

If all model assumptions are fulfilled, the motor noise model parameter *σ*^2^ will be linearly related to the error-variance *δ*^2^ and we should therefore be able to estimate *σ*^2^ from *δ*^2^. For each subject we computed *σ*^2^ by minimizing the squared error between the model task performance (eq. 13) and the experimentally determined task performance. A scatter plot of the resulting *σ*^2^ over the error-variance *δ*^2^ revealed an approximate linear relationship between *σ*^2^ and *δ*^2^. We then determined the proportionality factor *α* by linear-least squares regression of the model *σ*^2^ = *αδ*^2^ and used it to compute *σ*^2^ from *δ*^2^. The linear-least squares regression was done for each subject separately, using only the *σ*^2^ and *δ*^2^ values from all other subjects to avoid overfitting.

### Estimating the influence of model parameters on performance difference between expert and naïve groups

To estimate how much a single model parameter causes the experts’ gain in performance we computed the performance of the model for naive group parameters but with one parameter (horizon, motor noise, delay or motor penalty) changed to expert group values. We also performed the opposite procedure, replacing each parameter for each participants of the expert with those of the naïve group. Using the Bayesian inference approach described in the previous section, we replaced the full posterior distribution of the affected parameter with the posterior distribution from the other group. This procedure was carried out for each subject separately and the posterior of the affected parameter was replaced by the posterior of each subject from the other group separately. We then computed the posterior of the performance curve and from that the expected values of the performance by averaging. Hence, we obtained for each parameter change *N*_*e*_ ∙ *N*_*n*_ performance curves where *N*_*e*_ and *N*_*n*_ are the number of subjects in the expert and naïve group, respectively. These performance curves were averaged and compared to the average performances for the expert and naïve groups obtained for the fitted model (see Results for details).

Parts of the modelling computations were run on the high-performance computing cluster NEMO of the University of Freiburg (http://nemo.uni-freiburg.de) using Broadwell E5-2630v4 2.2 GHz CPUs.

All analysis code is available at https://github.com/dkobak/path-tracking.

## Results

### Learning the Tracking Skill

We designed an experiment where subjects had to a track a path moving towards them at a fixed speed (Fig. 1A and Methods). The narrow and wiggly path was moving downwards on a computer screen while the cursor had a fixed vertical position in the bottom of the screen and could only be moved left or right. Accuracy, our performance measure, was defined as the fraction of time that the cursor spent inside the path boundaries. One group of subjects (the expert group, N=32) trained this task for 30 minutes on each of 5 consecutive days. Another group (the naïve group, N=30) did not have any training at all. Both groups then performed a testing block that we describe below.

Over the course of five training days, the experts’ accuracy increased from 66.9±8.0% to 79.6±6.4% (mean±SD across subjects, first and last training day respectively) as shown on Figs 1B-C, with the difference being easily noticeable and statistically significant (p=8 ∙ 10^−7^, z=4.9, Wilcoxon signed rank test; Cohen’s d=1.8, N=32). As all paths generated during the training were different, this difference cannot be ascribed to memorizing the path, therefore this improvement represents the genuine acquisition of the skill of path tracking.

### Searchlight testing

To unravel the mechanisms of skill acquisition we designed testing trials called “searchlight trials”, during which subjects had to track curved paths as usual but could only see a certain part of the path (fixed distance s) ahead of the cursor. The searchlight length *s* varied between 10% and 100% of the whole path length in steps of 10% (the minimal s was ~3cm) to probe subjects’ planning horizon. Searchlight testing was conducted after 5 days of training for experts or immediately for novices. During the testing block all subjects completed 30 one-minute-long trials (three repetitions of each of the 10 values of s). The average accuracy at full searchlight *s*=100% was 82.8±7.5% for the expert group and 65.7±8.4% for the naïve group (mean±SD across subjects), with the difference being highly significant (p=2 ∙ 10^−9^, z=6.0, Wilcoxon-Mann-Whitney ranksum test, Cohen’s d=2.2, N=62). The performance of the naïve subjects matched the initial performance of the expert subjects on their first day of training.

Before we present the rest of the data, let us consider several possible ways in which the accuracy can depend on the searchlight length (Fig. 2A). For each subject, accuracy should be a non-decreasing function of searchlight length. The data presented in Poulton (1974) indicate that this function tends to become flat, i.e. subjects reach a performance plateau, after a certain value of the searchlight length that we will call *planning horizon* (Fig. 2A, top), while we assume all subjects will be constrained to the similar poor performance at the smallest searchlight. For the expert group, this function has to reach a higher point at s=100%, but it could do so because the initial rise becomes steeper, for example due to lower motor variability (Fig. 2A, bottom left), or because the initial rise continues longer, i.e. planning horizon increases (Fig. 2A, bottom right), or possibly both.

**Figure 2.**
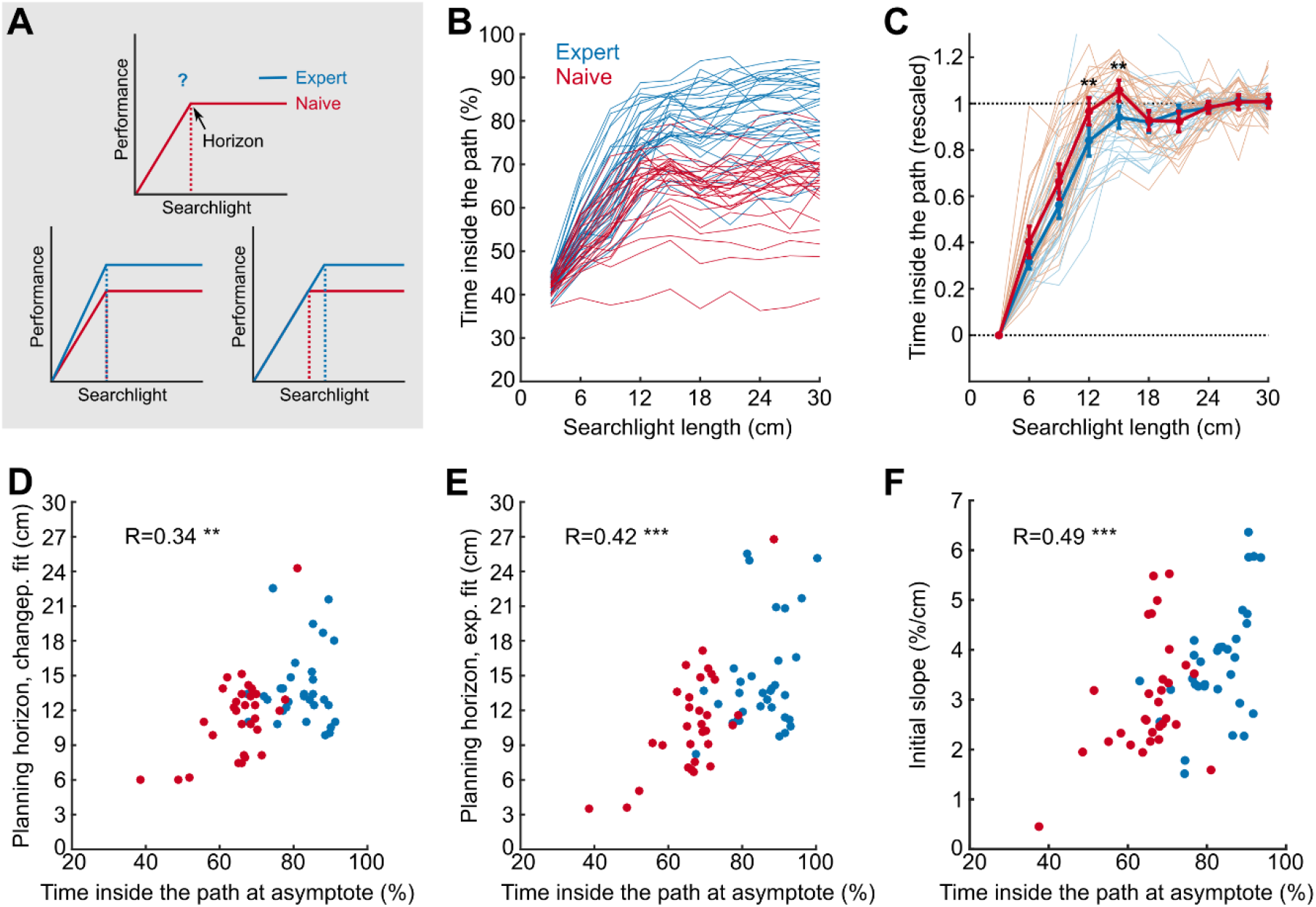
Searchlight testing. (A) Expert subjects were trained to have a higher performance at full searchlight length (top). This could be achieved by an increased initial slope (bottom left) at smaller searchlight length and/or an increased planning horizon as indicated with dashed vertical lines (bottom right). (B) Mean tracking performance for each searchlight length for each individual subject, in blue for the expert group and in red for the naïve group. Faint lines show individual subjects and bold lines show group means. (C) Mean tracking performance for each searchlight length, rescaled for each subject to start at 0 and end at 1 (see text). Error bars indicate 95% confidence intervals around the means, stars indicate significance between the groups (**: p<0.01, Wilcoxon rank sum text, Bonferroni-Holm corrected for multiple comparisons). (D-E) Planning horizon for each subject was defined by fitting a changepoint linear-constant curve (D) or an exponential curve (E) (see text). Both models yield an asymptote performance for each subject; the changepoint model yields a horizon length and the exponential fit yields an “effective” horizon length. The scatter plots show relation between the asymptote performance (as a proxy for subjects’ skill) and their planning horizon. Spearman’s correlation coefficients are shown on the plot (**: p<0.01, ***: p<0.001). Colour of the dot indicates the group. (F) Relationship between the asymptote performance and the initial slope in in the changepoint linear-constant model, colours and values as in D&E (***: p<0.001).

Fig. 2B shows subjects’ accuracy in the searchlights trials as a function of the searchlight length s. All subjects were strongly handicapped at short searchlights, and at the shortest searchlight the performance of the two groups was similar with experts being only marginally better (42.5±2.3% for the expert group, 41.4±1.8% for the naïve group, p=0.042, z=2.0 Wilcoxon ranksum test; Cohen’s d=0.5, N=62).

Visual inspection of Fig. 2B suggests that both effects sketched in Fig. 2A contribute to expert performance. (i) the planning horizon for the expert group was longer than for the naïve group; and (ii) the expert group had higher accuracies in the initial part of the performance curve, before the performance plateaus, which could be explained by decreased motor variability.

To better visualize the change in performance across searchlight lengths, we linearly rescaled each subject’s performance curve, first by subtracting the mean performance at s=10% and then by dividing by the asymptote performance (computed as the mean performance across s=80-100%). The resulting curves all start at 0 and end at 1 (Fig. 2C). We observed a significant difference between the groups at s=40% & 50% (p=0.005 and p=0.004 respectively, Wilcoxon ranksum test, p-values adjusted for testing 6 searchlight lengths between 20% and 70% with Holm-Bonferroni procedure, N=62), indicating that while naïve subjects had reached their plateau by then, the expert subjects kept increasing their performance. For this analysis we removed two naïve subjects with essentially flat searchlight curves (Fig. 1B), as rescaling those did not lead to meaningful results.

To investigate individual differences in tracking skill, we estimated the planning horizons of individual subjects (Fig. 2D). For this we fit each subject’s performance (*y*) with a changepoint linear-constant curve (see Methods), where the location of the changepoint defines the horizon length. We found that the novice group had an average horizon length of 11.5±3.6cm (mean±SD; median: 12.0cm) and the expert group a horizon length of 14.2±3.5cm (median: 13.2cm), with statistically significant difference (p=0.007, z=2.7, Wilcoxon ranksum test; Cohen’s d=0.8, N=62). We also found a positive correlation between the horizon length and the asymptotic performance (R=0.34, p=0.006, Spearman correlation, N=62).

In addition to the changepoint model, we also quantified the “effective” planning horizon using a single exponential to fit the individual subjects’ performance data (see Methods). This analysis confirmed our results (Fig. 2E). We again observed a significant difference in the effective horizon length between the two groups (14.76±4.6cm vs. 11.04±4.7cm, means±SD for both groups, medians: 13.6cm and 10.7cm, p=0.002, z=3.0, Wilcoxon ranksum test; Cohen’s d=0.8, N=62). Again, we found a positive correlation between the asymptote performance and the effective horizon length (R=0.43, p=0.0008, Spearman correlation, N=62).

Not only was planning horizon positively correlated with tracking skill (the asymptote accuracy), but also the initial slope of the changepoint model (3.7±1.2 %/cm vs. 3.0±1.2 %/cm, mean±SD; medians: 3.6 %/cm vs. 2.6 %/cm). Fig. 2F shows that there was a positive correlation between the initial slope and asymptote accuracy (R=0.49, p=6.10^−5^ Spearman correlation, N=62) as well as a clear difference in the initial slope between the groups (p=0.008, z=2.6, Wilcoxon ranksum test; Cohen’s d=0.6, N=62).

We therefore conclude that the difference between expert and naïve performances is a combination of both possibilities presented in Fig. 2A. Using the expert and naive median estimates of the intercept, the slope, and the horizon in the changepoint model, we can estimate the contribution of both effects on the asymptote performance. The changepoint model asymptote performance for the naive group was 63.5%, compared to 78.7% for the expert group. The model performance of the expert group at the naive horizon was 74.2%. Hence, approximately 71% of the expert performance gain of 15.2%, was due to the increase in the initial slope (possibly due to lower motor variability), and the remaining 29% can be attributed to the increase in planning horizon. The identical procedure with mean model parameter estimates instead of median estimates, yields 44% attributable to motor acuity and 56% attributable to planning horizon. We conclude that between a third and a half of the expert performance gain is attributable to their increase in planning horizon.

### Trajectory analysis

Naïve subjects performed worse than the expert subjects at long searchlights but all subjects performed almost equally badly at short searchlights. What kinematic features can these differences be attributed to?

Clearly, at short searchlights, performance has to be reactive. To measure how quickly changes in the path were reflected in the motor commands, we computed the time lag between cursor trajectory and path midline (the lag maximizing cross-correlation between them). As Fig. 3A shows the lag was ~200 ms at s=10% for all subjects and dropped to ~0 ms at s=50% for the expert group. While many naïve subjects also decreased their lags to zero, 10 out of 30 never achieved the 0 ms lag. The five naïve subjects showing the largest lags at large searchlights were also those with the worst performance (Fig. 3B). Therefore, there was a strong negative correlation between the asymptote lag (mean across s=80-100%) and the asymptote performance (mean across s=80-100%) of R=-0.58 (Fig. 3B, p=8 ∙ 10^−7^, Spearman correlation, N=62).

**Figure 3.**
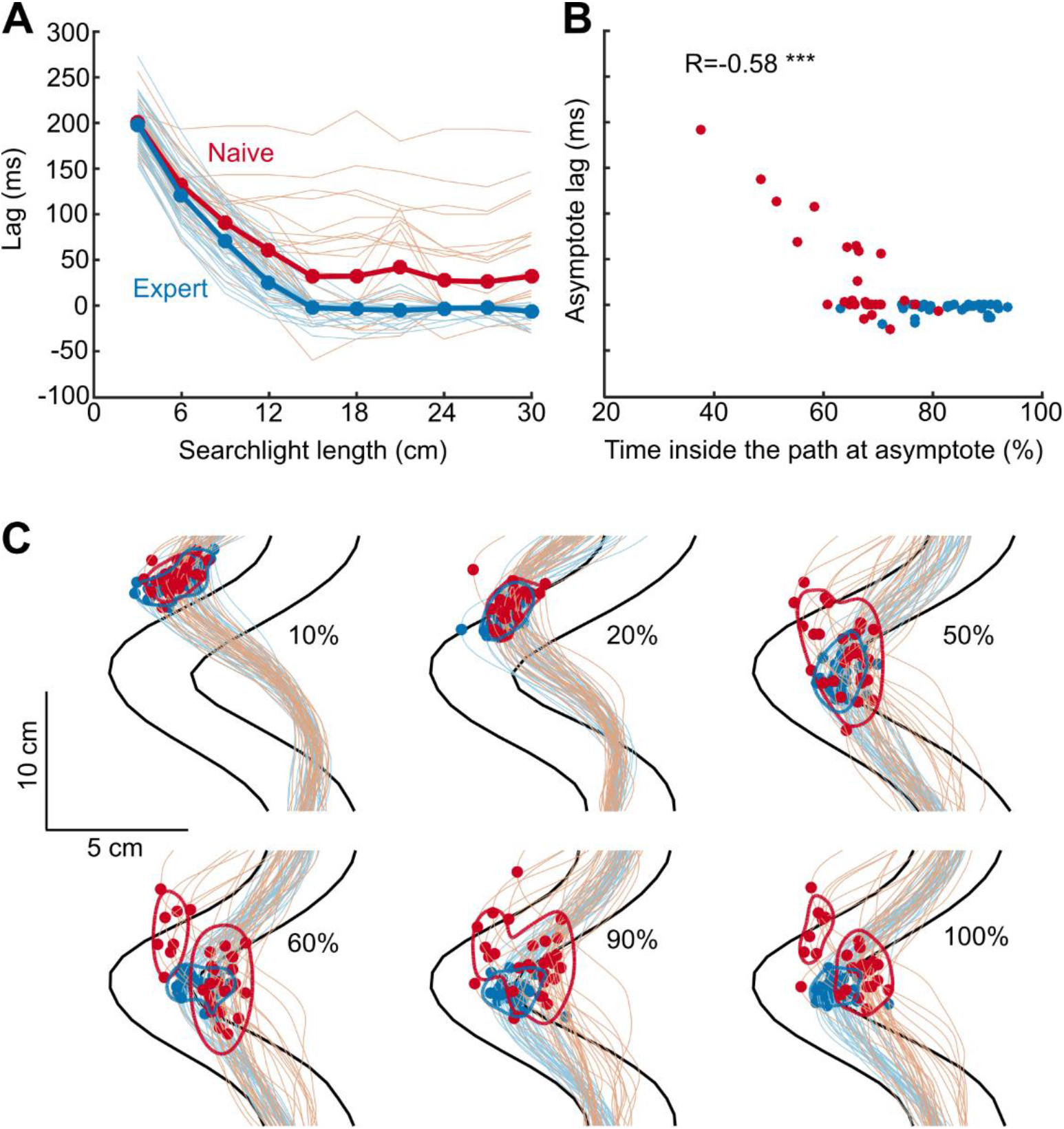
Analysis of trajectories. (A) Mean time lag between cursor trajectory and path midline, for each searchlight length for each individual subject (faint lines) and mean of per-subject values (bold lines), in blue for the expert group and in red for the naïve group. (B) Asymptote lag and asymptote performance across subjects. Correlation coefficient is shown on the plot (***p<0.001). Colour of the dot indicates the group. (C) Average per-subject trajectories in sharp bends (leftward bends were flipped to align them with the rightward bends). Each trajectory is averaged across approximately 40 bends (the number of bends varied across searchlight lengths). Colour of the lines indicates the group. Black lines show average path contour. Dots show turning points of the trajectory. Contour lines show the kernel density estimate 75% coverage areas. Subplots correspond to searchlight lengths s=10%, 20%, 50%, 60%, 90% and 100%.

Next, for each testing path we found all segments exhibiting sharp leftward or rightward bends (see materials and methods, our inclusion criteria yielded 13±5 segments per path, mean±SD). For each searchlight length s and for each subject, we computed the average cursor trajectory over all segments (N=38±8 segments per searchlight) after aligning all segments on the bend position (Fig. 3C, leftward bends were flipped to align them with the rightward bends). At s=10% all subjects from both groups follow very similar lagged trajectories, resulting in low accuracy. As searchlight increases, expert subjects reach zero lag and choose more and more similar trajectories, whereas naïve subjects demonstrate a wide variety of trajectories with some of them failing to reach zero lag and others failing to keep the average trajectory inside the path boundaries. To visualize this, we plotted the kernel density estimate 75% coverage contour of inflection points for each group. As the searchlight increases, the groups become less overlapping and the naïve group appears to form a bimodal distribution (Fig. 3C).

In summary, at very short searchlights all subjects performed poorly because in this reactive regime their trajectories lagged behind the path. At longer searchlights the expert subjects were able to plan their movement to accommodate the bends (the longer the searchlight the better), but naïve subjects failed to do so in various respects: either still lagging behind or not being able to plan a good trajectory.

### Receding horizon model analysis

Next, we modelled subjects’ behaviour by receding horizon control (RHC) to illustrate that such an approach is able to capture some crucial features of the behavioural data. In RHC a sequence of motor commands is computed to minimize the expected cost over a future time interval of finite length, i.e. the horizon. After the first motor command is applied, the optimization procedure is repeated using a time interval shifted one time step ahead. See Methods section for a more detailed and formal description of RHC. As cost function, we used the weighted sum of a measure of inaccuracy (i.e. probability of being outside the path) and the magnitude of the motor cost (see Methods for details). Cost function with a similar trade-off between movement accuracy and motor command magnitude have been used previously to describe human motor behaviour in different tasks (Braun et al. 2009; Diedrichsen 2007; Todorov and Jordan 2002). The model has four different parameters: horizon (*h*), motor noise (*σ*^2^), motor delay (*τ*) and motor command penalty weight (*λ*.

We ran the model on the experimental paths to obtain simulated movement trajectories from which task performance and lag could be computed in the same way as for the experimental trajectories (Fig. 2 and 3). Our simulations revealed that both, a larger model horizon as well as a smaller motor noise parameter increased the task performance and decreased the lag (Fig. 4). Hence, the experimentally observed higher performance and smaller lag of expert subjects compared to naive (Fig. 2B and 3A) could be explained either by an increased model horizon or by reduced motor noise in the model. However, the searchlight length at which the task performance of the model reached a plateau increased with model horizon and did not change or even decreased with a smaller motor noise parameter (Fig. 4A, C). Experimentally, on the other hand, we observed that subjects with a higher task performance reached their performance plateau at higher searchlights (Fig. 2D, E). This correlation between performance and plateau onset, that was observed experimentally, cannot be explained by the variation of the motor noise parameter across subjects, but is only consistent with an increase of the model planning horizon for subjects with higher performance.

**Figure 4:**
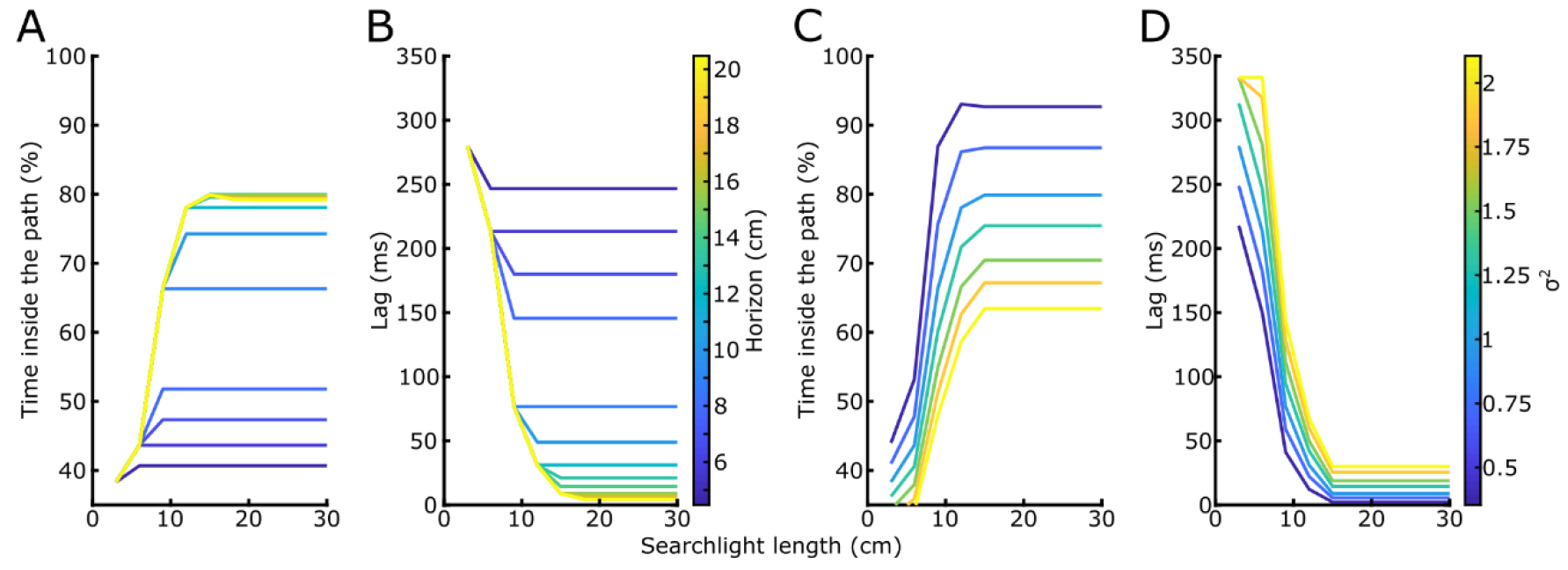
Task performance and lag as a function of searchlight length for model simulations with different horizons (A,B) or different amounts of motor noise (C,D). A motor noise of *σ*^2^=1 was used for (A,B) and a horizon of *h*=15cm for (C,D). The motor delay and motor command penalty weight were fixed at *τ*=200ms and *λ*=0.5 in all simulations.

Next, we used Bayesian inference to estimate the model parameters from the experimentally observed movement trajectories (see Methods for details). Based on inferred distributions of parameter values, we then predicted task performance and lag for each subject. To avoid over-fitting cross-validation was used, i.e. fitting and prediction was done on different trials. Model task performance and lag resembled the experimentally observed task performance and lag with regard to their change across searchlights as well as with regard to the difference between naïve and the expert subjects (Fig. 5A,B). On a single subject and trial level there was a high correlation between model and experimental task performance (Fig. 5C, Spearman correlation r=0.9, R^2^=0.84) and lags (Fig. 5D, Spearman correlation r=0.87, R^2^=0.88).

**Figure 5:**
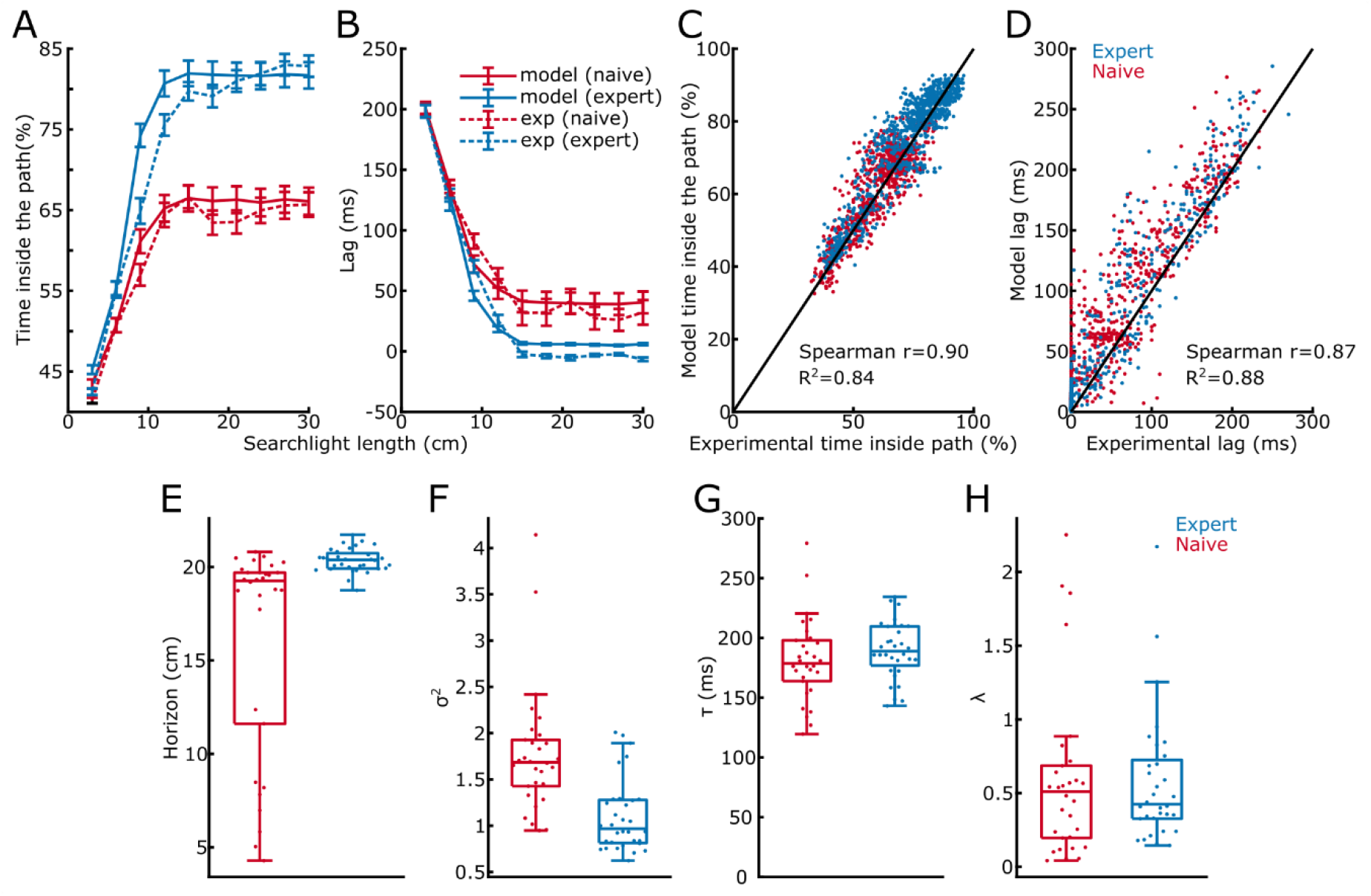
Comparison between the receding horizon model and subjects’ behaviour. A,B: Task performance and lag as a function of the searchlight for expert and naïve subjects for the experiments and model simulations. C,D: Scatter plot of model and experimental task performance and lag for each trial of each subject. E-H: Model parameters for the subjects from the naïve and the expert group. Each dot depicts one subject, boxplots show medians as well as first and third quartiles.

We compared the estimated model parameters between expert and naïve subjects. The fitted model horizon was higher for the expert group than for the naïve group (Fig. 5E, Wilcoxon ranksum test: z=4.84, p=1·10^−6^, N=62) and was correlated with the horizon obtained from the change point analysis (Spearman correlation, r=0.48, p=7·10^−5^, N=62) and the exponential fits (Spearman correlation, r=0.43, p=6·10^−4^, N=62). One caveat here is that there was large uncertainty in the estimates of model horizon for most subjects, and the exact values shown in Fig. 5E might be systematically biased due to some model misspecification (in particular, note that the values in Fig. 5E are all larger than the estimates in Fig. 2D,E). That said, the RHC model estimates qualitatively agree with our earlier estimates that the expert horizons were larger than the naive horizons.

The fitted motor noise was significantly lower for the expert than for the naïve group (Fig. 5F; Wilcoxon ranksum test: z=4.66, p=3·10^−6^, N=62) while the delay and the penalty parameters were not different (Fig. 5G,H; delay: Wilcoxon rank sum test, z=1.50, p=0.13; penalty: Wilcoxon rank sum test, z=0.528, p=0.60, N=62). In the model, lower motor noise lead to steeper initial accuracy slope (Fig. 4C). The expert group having lower estimated motor noise hence agrees well with our observation that experts had steeper initial accuracy slope (Fig. 2F).

Using the model fits obtained above, we estimated how much of the experts’ gain in asymptote performance was due to increased horizon vs. decreased noise. To do this, we simulate the model with naive group parameters but expert group horizons (see Methods). This brings the performance almost half-way to the expert performance (for large searchlights the performance levelled off at 72% instead of 82% with lower horizon, compared to 66% for the naïve subjects). We observe roughly the same increase (to 75%) when we simulate the model with naive group parameters but expert group noise levels. Similarly, when we use expert group parameters but naive group horizons or noise levels, the performance drops approximately half-way to the naïve accuracy (74% for naïve horizon, 71% for naïve motor noise). In contrast, the delay and the motor penalty parameters had less influence on the asymptote performance (63% and 64% for naïve group parameters with expert delay or motor penalty; 80% for expert group parameters with naïve delay or motor penalty). From this we conclude that the increase in the experts’ performance was caused by equal measures through an increase in planning horizon and the decrease in motor noise. This is in a good qualitative agreement with the conclusions we presented earlier based on the linear changepoint fits.

## Discussion

We used a paradigm that allowed us to study skill development when humans had to track an unpredictable spatial path. The skill requires fast reactions to new upcoming bends in the road, but also a substantial “planning ahead” component – i.e. the anticipation and preplanning of movements that have to be made in the near future. We used the accuracy, i.e. the fraction of time the cursor was inside the path boundaries, as the measure of performance. We observed a substantial improvement in accuracy after 5 days of training (Fig. 1B,C). The paths were different on every trial, so the improvement in performance cannot be attributed to a memory for the sequence.

What changes in the motor system occur through learning that allowed skilled subjects to perform better? One component of this improvement has been previously called “motor acuity” (Shmuelof et al. 2012, 2014) and corresponds to the subjects’ ability to execute motor commands more accurately, i.e. due to the lower motor variability. We hypothesized that an additional component is an increased ability to take into account approaching path bends and to prepare for an upcoming movement segment. We directly estimated both effects by using a searchlight testing where only a part of the approaching curve was visible. In agreement with our hypothesis, we found that subjects with a higher tracking skill demonstrated larger planning horizons: on average ~14cm for the expert group vs. ~11cm for the naïve group, corresponding to the time horizons of ~0.4s and ~0.3s respectively. Our results suggest that the increase in planning horizon is not an epiphenomenon but is causally related to the performance increase, as expert subjects showed worse performance when the searchlight was reduced below their planning horizon (Figure 2C). We estimated that in our experiments between a third and a half of the increased performance after practice can be attributed to an increased planning horizon while the rest can be accounted for by a reduction in the motor variability which may be interpreted as higher motor acuity.

The expert group showed higher initial slope of the searchlight-accuracy curve. We interpreted this as an indirect evidence for lower motor variability, even though other explanations for higher slope are in principle also possible. Our assumption was that as long as the searchlight lengths does not exceed a subject’s horizon, all subjects (expert and naive) are able to use information about the whole visible path chunk. The results of the RHC fits showed a clear difference in motor noise between the groups, in agreement with our interpretation that the expert group had lower motor variability.

Note that “planning”/“preparing” the movement can be interpreted differently depending on the computational approach. In the framework of optimal control (Todorov and Jordan 2002), subjects do not plan the actual trajectory to be followed, but instead use an optimal time-dependent feedback policy and then execute the movement according to this policy. The observed increase in planning horizon can be interpreted in the framework of model predictive control, also known as receding horizon control, RHC (Kwon and Han 2005). In RHC, the optimal control policy is computed for a finite and limited planning horizon, which may not capture the whole duration of the trial. This policy is then applied for the next control step, which is typically very short, and the planning horizon is then shifted one step forward to compute a new policy. Hence, RHC does not use a pre-computed policy, optimal for an infinite horizon, but a policy which is only optimal for the current planning horizon. Increasing the length of the planning horizon is therefore likely to increase the accuracy of the control policy. In our experiments this would allow for a larger fraction of time spent within the path boundaries. We designed a simple RHC model to test directly which components in the model would have to change through training to quantitatively explain the subject’s behaviour. The dynamics of movement and the cost function were modelled in line with previous studies that used optimal control to describe human behaviour in various motor control and learning tasks (Braun et al. 2009; Diedrichsen 2007; Todorov and Jordan 2002). We fitted the RHC model to the behaviour of each subject and found that it was able to fit the data very accurately (Fig. 5). The experimentally observed differences between expert and naïve subjects were reflected in the model fits by higher planning horizons and lower motor noise parameters in the expert group. Our findings, thus, demonstrate that subjects’ behaviour can be understood in the context of RHC, and longer planning horizons of the expert group indicate that subjects learn how to take advantage of future path information to improve motor performance.

Despite a clear difference in the distribution of planning horizons between the naive and the expert groups (Fig. 2D), there was a substantial overlap: the planning horizon of many naive and expert subjects were similar. While this might simply reflect a moderate effect size combined with inter-subject variability and measurement noise, it also remains a possibility that the difference between groups was largely caused by those naive subjects with very low horizons and expert subjects with very high horizons.

### Related work

Ideas like the RHC were put forward in a recent study (Ramkumar et al. 2016) that suggested that movements are broken up in ‘chunks’ in order to deal with the computational complexity of planning over long horizons. That study suggests that monkeys increase the length of their movement chunks during extended motor learning over the course of many days which may be explained by monkeys increasing their planning horizon with learning. At the same time, the efficiency of movement control within the chunks improved with learning which may also be the result of a longer horizon. Despite these potential consistencies with our approach we note that in their model Ramkumar et al. (2016) assumed that ‘chunks’ are separated by halting points (i.e. points of zero speed) and movements within ‘chunks’ are optimized independently from each other. Our RHC model does not have independent movement elements but movements are optimized continuously.

Even though our study, to the best of our knowledge, is the first to directly investigate the evolution of the planning horizon during continuous path tracking, an increase in the planning horizon after learning has been recently demonstrated when learning sequences of finger movements (Ariani et al. 2020). Similar path tracking tasks have been used before (Poulton 1974). Using a track that was drawn on a rotating paper roll, these early studies found that the accuracy of the tracking increased with practice and with increasing searchlight length (which was modified by physically occluding part of the paper roll, (Poulton 1974), p 187). These studies, however, did not investigate the effect of learning on the planning horizon.

More recent studies used path tracking tasks where the goal was to move as fast as possible while maintaining the accuracy (instead of moving at a fixed speed). In all of these studies the identical path was repeatedly presented. In one study subjects had to track a fixed maze without visual feedback and learnt to do it faster as the experiment progressed (Petersen et al. 1998); there the subjects had to once “discover” and then remember the correct way through the maze. In another series of experiments, Shmuelof et al. asked subjects to track two fixed semi-circular paths. Subjects became faster and more accurate over the course of several days (Shmuelof et al. 2012), but this increase in the speed and accuracy did not generalize to untrained paths (Shmuelof et al. 2014). In contrast to these previous path tracking studies, we used randomly generated paths throughout the experiment. By investigating the generalization of the path tracking skill to novel paths we could reveal an increasing planning horizon with learning.

## Conclusion

In conclusion, we have established that people are able to learn the skill of path tracking and improve their skill over 5 days of training. This increase in motor skill is associated with the increased motor acuity and increased planning horizon. The dynamics of preplanning can be well described by a receding horizon control model.

## Acknowledgements

The study was in parts supported by the German Federal Ministry of Education and Research (BMBF) grant 01GQ0830 to BFNT Freiburg-Tübingen. The authors acknowledge support by the state of Baden-Württemberg through bwHPC and the German Research Foundation (DFG) through grant no INST 39/963-1 FUGG. The authors also thank the ‘Struktur-und Innovationsfonds Baden-Württemberg (SI-BW)” of the state of Baden-Württemberg for funding.

## Author Contribution

Conceptualization, LB. DK. and CM; Methodology, LB. DK. JD and CM; Formal Analysis, LB. DK and CM; Writing – Original Draft, LB. DK and CM; Writing – Review and Editing, LB, DK, JD and CM.

